# Affective contagion: how attitudes expressed by others influence our perception of actions

**DOI:** 10.1101/2021.05.05.442710

**Authors:** G. Di Cesare, A. Pelosi, S. Aresta, G. Lombardi, A. Sciutti

## Abstract

Vitality forms represent a fundamental aspect of social interactions, characterizing *how* actions are performed and *how* words are pronounced, on the basis of the agent’s attitude. The same action, such as a handshake, has a different impact on the receiver when it is performed kindly or vigorously, and the same happens with a gentle or rude tone of voice. In the present study, we carried out two experiments which aimed to investigate whether and how vocal requests conveying different vitality forms can influence the perception of goal-directed actions and to measure the duration of this effect over time. More specifically, participants listened to an actor voice pronouncing “give me” in a rude or gentle way, and then they were asked to observe the initial part of a rude or gentle passing action, continue it mentally and estimate its conclusion. Results showed that the perception of different vitality forms expressed by vocal requests influenced the estimated action duration. Moreover, we found that this effect was limited to a certain time interval, after which it started to decay.

## 1. Introduction

The observation of goal-directed actions performed by another individual allows one to understand what that individual is doing, why, and how he/she is doing it. During social interactions, observing how actions are performed or listening to the tone of voice, people are able to understand the internal state of others. Indeed, actions and speech dynamics represent fundamental aspects of social communication, defined “*vitality forms*” by Daniel Stern.^1^ Vitality forms characterize human behavior enhancing the quality of interactions. In particular, the expression of vitality forms enables the agent to communicate his/her mood while the perception of vitality forms allows the receiver to capture immediately the attitude of the agent.^2^ For example, a hand gesture can be performed vigorously or kindly, a tone of voice can be unpleasant or pleasant, suggesting that the agent has a negative or positive attitude towards the receiver.

Different fMRI studies have investigated the neural correlates involved in the processing of vitality forms, showing that the dorso-central insula has a crucial role in the perception and execution of actions conveying gentle and rude vitality forms.^3-6^ Moreover, the same authors found that this insular sector was also activated during the processing of action words, action sounds and touches conveying different vitality forms.^7-9^ Considering the multimodal role of the dorso-central insula, it is plausible to hypothesize that, during interpersonal relations, the insula encodes vitality forms of speech, actions and touch and automatically transforms them into a motor domain, allowing the receiver to understand the affective state of the other and prepare an appropriate motor response.^6^ This hypothesis is corroborated by a kinematic study,^10^ which showed that requests conveying vitality forms (rude/gentle) presented in visual modality, auditory modality or mixed (visual and auditory), influenced the execution of actions performed by the receiver. These findings represent first evidence that vitality forms expressed by an agent affect the motor behavior of the receiver, opening new future research question that need to be addressed. An interesting issue is to understand whether the encoding of vitality forms expressed by an agent may influence the perception of the receiver. For this purpose, the present study aims to investigate two points: 1) how gentle and rude vitality forms may affect the way of perceiving actions during an action observation task; 2) quantify the duration of this effect. In this respect, we carried out two behavioural experiments. In the first experiment, participants were required to perform a cognitive task. Specifically, they listened a voice of an actor/actress pronouncing “give me” (“dammi”: Italian verb) in a rude or gentle way. After listening the actor/actress vocal request, participants observed video clips showing only the initial part of a rude or gentle action (passing an object) and were required to continue mentally the action indicating its end. Taking advantage of the task used for this first experiment, in the second experiment, between the vocal request and the action presentation, time delays of different durations were added. Concerning the first experiment, we hypothesized an influence of different vitality forms conveyed by vocal requests on the estimation of action duration. Since vitality forms are one of the first information we gather from others, concerning the second experiment we hypothesized that this effect was limited to a specific time interval, after which it decayed.

## 2. Material and methods

### First Experiment

#### 2.1 Participants

Thirty healthy right-handed volunteer subjects (20 females and 10 males, mean age = 24.4; SD = 2.87) participated in this study. All participants had normal or corrected to normal visual acuity. Nobody reported a history of psychiatric or neurological disorders or current use of psychoactive drugs.

#### 2.2 Visual Stimuli

Participants sat comfortably in front of a table on which a laptop was placed, positioning their right hand on the mouse. The stimuli consisted of video clips showing an actor/actress passing different objects (a packet of crackers, a ball, a bottle, a cup) to another person, in a gentle or rude way. It is important to note that, in order to facilitate the motor representation of the observed action in participants, video-clips showed actions performed by a right hand with an egocentric perspective. More specifically, videos were obscured so that participants observed only a portion of the entire duration (Figure 1). For rude actions, stimuli presented could last 200ms, 250ms, 300ms or 350ms, corresponding respectively to 28%, 35%, 42% and 50% of the total duration (700ms). For gentle actions, stimuli presented could last 340ms, 420ms, 500ms, or 600ms, corresponding respectively to 28%, 35%, 42% and 50% of the total duration (1200ms). Stimuli have been created using E-Prime software and were presented in a random order.

**Figure 1.**
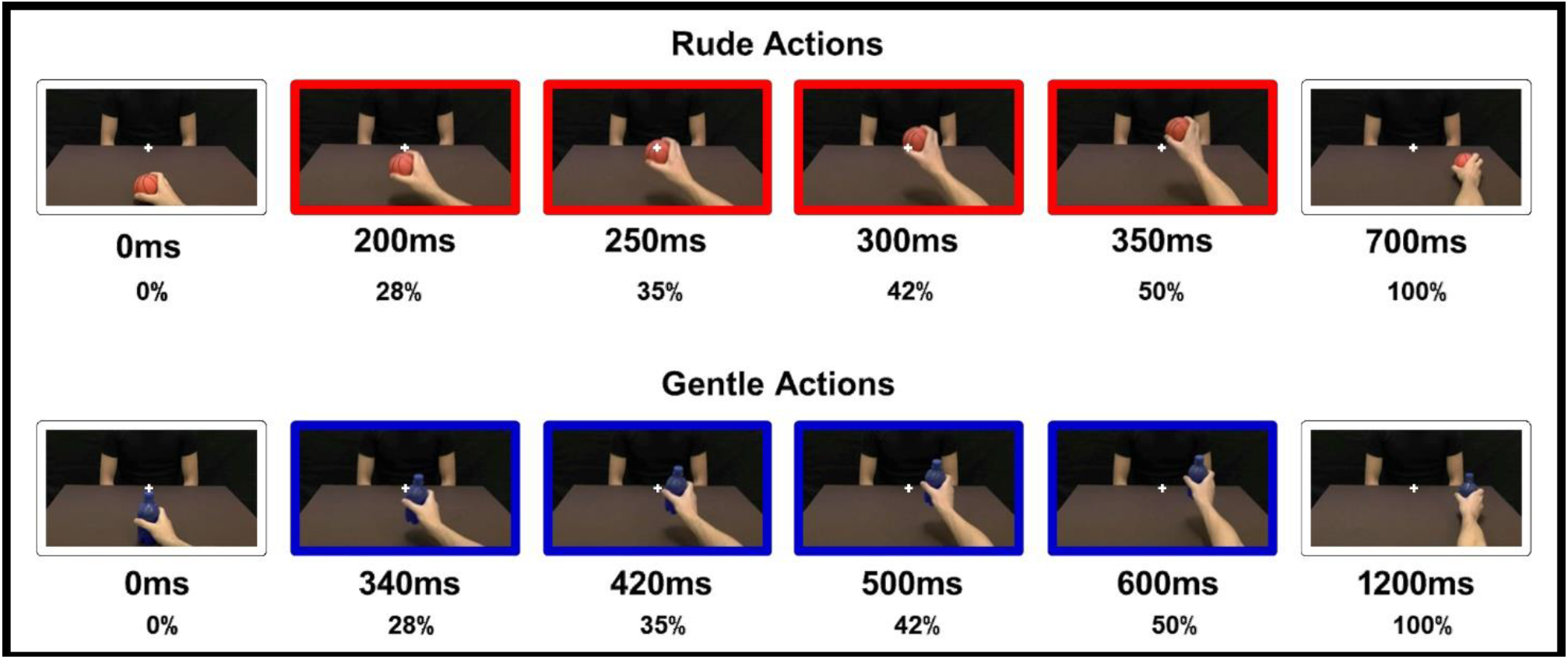
Video stimuli presented to participants. For rude actions (red colour): 200ms, 250ms, 300ms or 350ms, corresponding respectively to 28%, 35%, 42% and 50% of the total duration (700ms). For gentle actions (blue colour): 340ms, 420ms, 500ms or 600ms, corresponding respectively to 28%, 35%, 42% and 50% of the total duration (1200ms).

#### 2.3 Task and Experimental Paradigm

During the experiment participants were stimulated with a vocal request, consisting in a voice of an actor/actress, pronouncing “give me” (Italian verb: “dammi”) in a rude or gentle way. After each vocal request, participants were presented with video stimuli described above and were required to continue mentally the action partially observed and to estimate its end, pressing the mouse. Each vocal request was recorded using a condenser microphone (RODE NT1) placed 30 cm in front of the actors and digitized with a phantom powered A/D converter module (M-AUDIO M-TRACK). After recording, the audio files were processed with COOL EDIT PRO software to obtain the final version of the stimuli. Rude and gentle vocal requests differed for parameters such as the wave amplitude (Figure 2AC) and the pitch (Figure 2BD).

**Figure 2.**
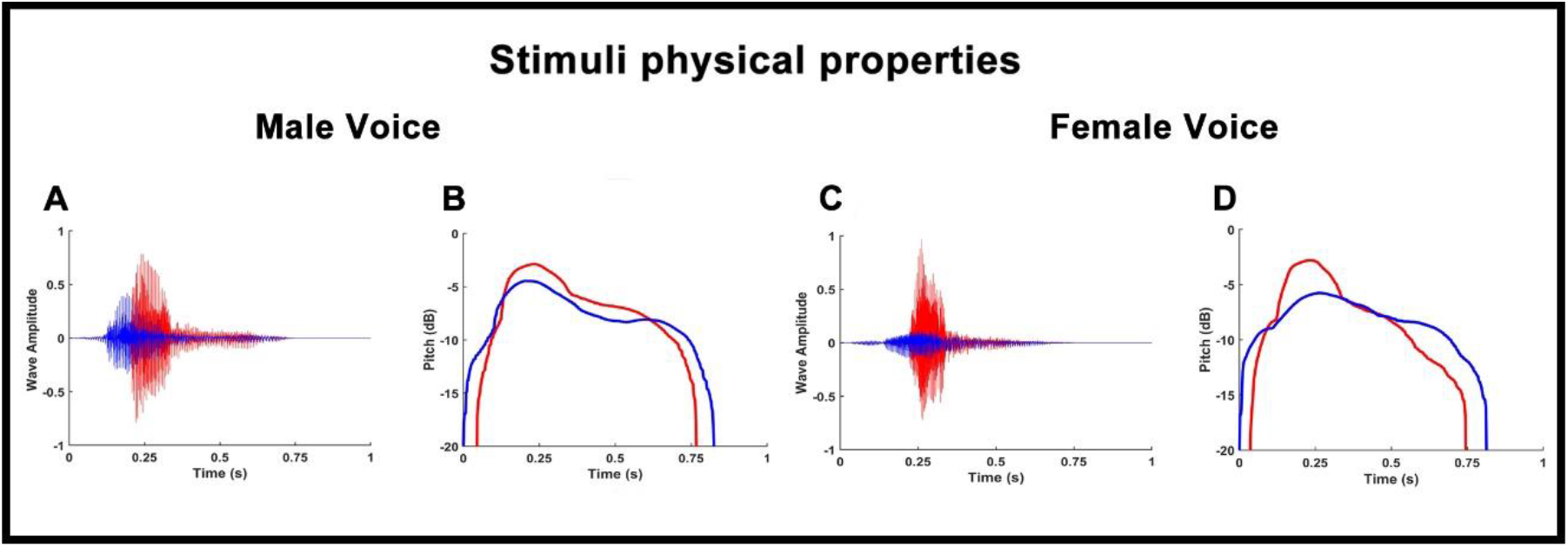
Graphs show the wave amplitude of male (A) and female (C) voices; the pitch of male (B) and female (D) voices. Red colour refers to rude vitality form. Blue colour refers to gentle vitality form.

Four conditions were randomly presented using E-prime software: 1) RDV_RDA (congruent condition): the vocal request and the observed action were both rude 2) GTV_GTA (congruent condition): the vocal request and the observed action were both gentle; 3) RDV_GTA (incongruent condition): the vocal request was rude and the observed action was gentle; 4) GTV_RDA (incongruent condition): the vocal request was gentle and the observed action was rude. Each video was presented seven times for each condition. Before the beginning of the experiment, participants were asked to perform a training session to assess recognition of audio and video stimuli and to become familiar with the experimental task. The experiment was composed of three different runs (Figure 3). In the first run, participants simply performed the task, without receiving vocal requests before, to assess their capacity to correctly estimate the duration of rude and gentle actions. This run was used as a baseline and lasted 2 minutes. In the second and third runs, participants listened to the vocal request, expressed with rude or gentle vitality form, and then they observed the beginning of the action and estimated its conclusion. Half of the vocal requests were performed by an actress and the other half by an actor. The duration of the second and third runs was 9 minutes.

**Figure 3.**
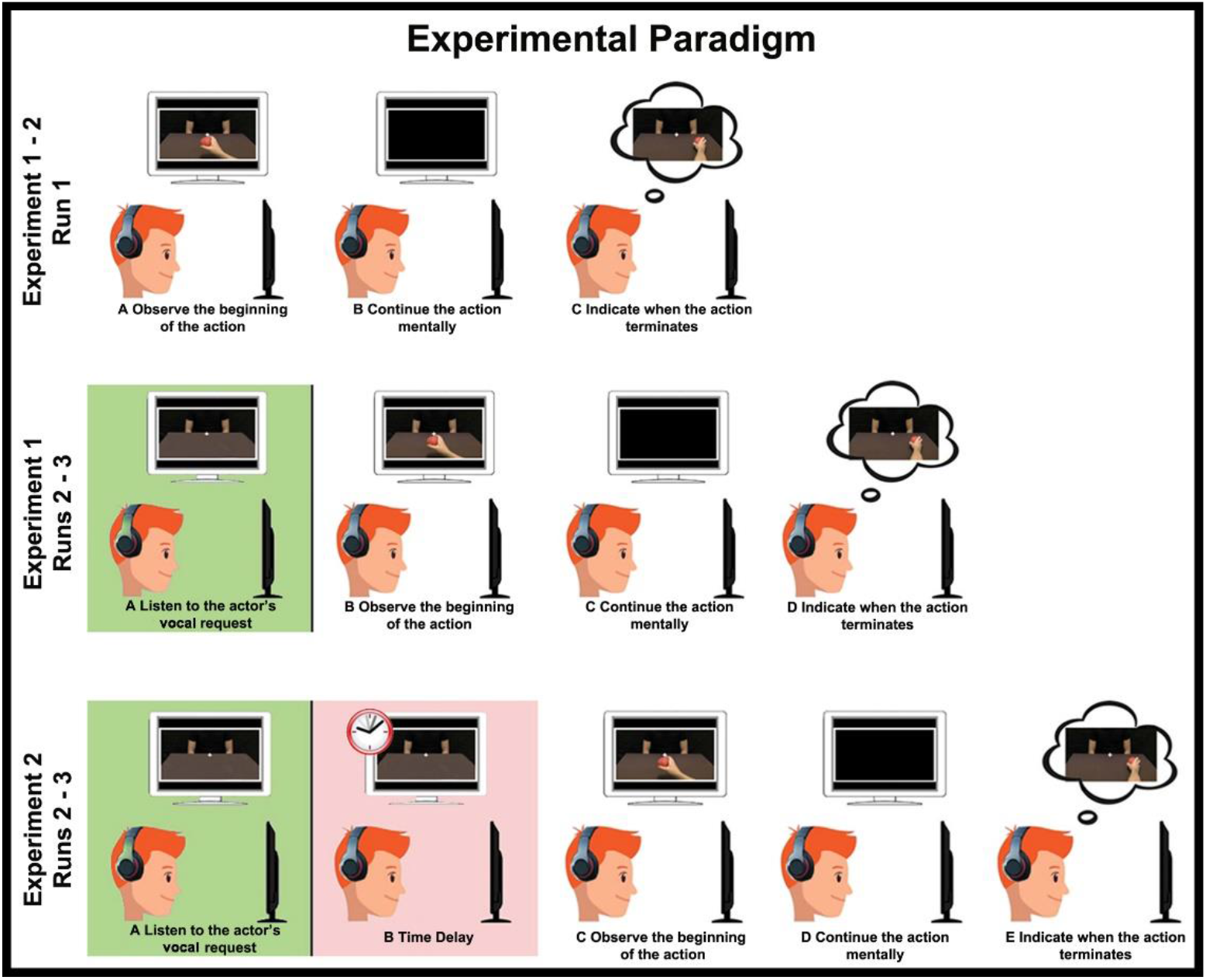
For both the experiments, in run1: A) participants observed the beginning of the action; B) continued the action mentally and C) estimated its conclusion. For the first experiment, in run2 and run3: A) participants listened to a rude/gentle vocal request B) observed the beginning of the action; C) continued the action mentally; D) estimated its conclusion. In the second experiment, run2 and run3 were the same, but between the vocal request and the video stimuli a time delay was inserted

### Second Experiment

#### 2.4 Participants

Thirty-eight healthy right-handed volunteer participants (21 females and 17 males: mean age = 24.4; SD = 4) took part in this study. All participants had normal or corrected to normal visual acuity. Nobody reported a history of psychiatric or neurological disorders or current use of psychoactive drugs.

#### 2.5 Task and Experimental Paradigm

From stimuli described above, in the second experiment participants were only presented with video clips of 200ms for rude actions and 340ms for gentle actions. The task they were asked to perform was the same of the first experiment: they listened to the vocal request expressed gently or rudely by the actor/actress, then they observed the initial part of the action and estimated its conclusion pressing the mouse. In order to estimate the duration of the effect of vocal request conveying vitality forms on the task performed after, time delays have been added between the vocal request and the presentation of video-clips. Time delays were 0ms (meaning no delay between vocal request and video stimuli), 400ms, 800ms, 1200ms or 1600ms. As in the first experiment, the second experiment was composed of three runs and stimuli were presented randomly using E-prime software (Figure 3).

## 3. Results

### 3.1 First Experiment

In order to evaluate the effect of vocal requests conveying different vitality forms on the perception of observed actions, we analyzed the participants’ responses (estimated action durations). Participants’ responses were modelled using two Repeated Measured GLM (general linear models). The significance level was fixed at p = 0.05. Before performing statistical analysis, the sphericity of data was verified (Mauchly’s test, p > 0.05) and the Greenhouse–Geisser correction was applied in case of sphericity violation (p < 0.05). The first GLM model comprised the participant’s response time related to videoclips of four different durations (340ms, 420ms, 500ms, 600ms) showing gentle actions in three experimental conditions (baseline: NOV_GTA, congruent: GTV_GTA, incongruent: RDV_GTA). The second GLM model comprised the participant’s response time related to videoclips of four different durations (200ms, 250ms, 300ms, 350ms) showing rude actions in three experimental conditions (baseline: NOV_RDA, congruent: RDV_RDA, incongruent: GTV_RDA). Results of the first GLM analysis indicated a significant difference among experimental conditions (F = 11.67, p < 0.01, partial-η^2^ = 0.28, δ = 0.99), durations (F = 2.76, p < 0.05, partial-η^2^ = 0.087, δ = 0.65) and an interaction between experimental conditions * durations (F = 2.39, p < 0.05, partial-η^2^ = 0.076, δ = 0.67). For each duration, post hoc analysis revealed a significant difference among experimental conditions (340ms: NOV_GTA vs RDV_GTA p < 0.001, GTV_GTA vs RDV_GTA p < 0.001; 420ms: NOV_GTA vs RDV_GTA p < 0.01, GTV_GTA vs RDV_GTA p < 0.001; 500ms: NOV_GTA vs RDV_GTA p < 0.05; 600ms: NOV_GTA vs RDV_GTA p < 0.001, GTV_GTA vs RDV_GTA p < 0.001; Bonferroni correction, see Figure 4A). Results of the second GLM analysis indicated a significant difference among experimental conditions (F = 5.39, p < 0.01, partial-η^2^ = 0.15, δ = 0.75), durations (F = 14.78, p < 0.001, partial-η^2^ = 0.33, δ = 1) and an interaction between experimental conditions * durations (F = 2.54, p < 0.05, partial-η^2^ = 0.081, δ = 0.73). For each duration, post hoc analysis revealed a significant difference among experimental conditions (200ms: NOV_RDA vs RDV_RDA p < 0.001, GTV_RDA vs RDV_GTA p < 0.001; 300ms: NOV_RDA vs GTV_RDA p < 0.001, RDV_RDA vs GTV_RDA p < 0.001; 350ms: NOV_RDA vs GTV_RDA p < 0.01, RDV_RDA vs GTV_RDA p < 0.0; Bonferroni correction, see Figure 4B). In order to quantify the effect of vocal requests on the action estimation for each duration, we compared values obtained in congruent conditions with those obtained in incongruent conditions [(RDV_GTA – GTV_GTA)*100 / GTV_GTA for gentle actions; (GTV_RDA – RDV_RDA)*100 / RDV_RDA for rude actions]. Results of this analysis revealed that the effect of vocal request on estimated action duration was consistent across different durations and for both gentle and rude actions (Figure 4CD).

**Figure 4.**
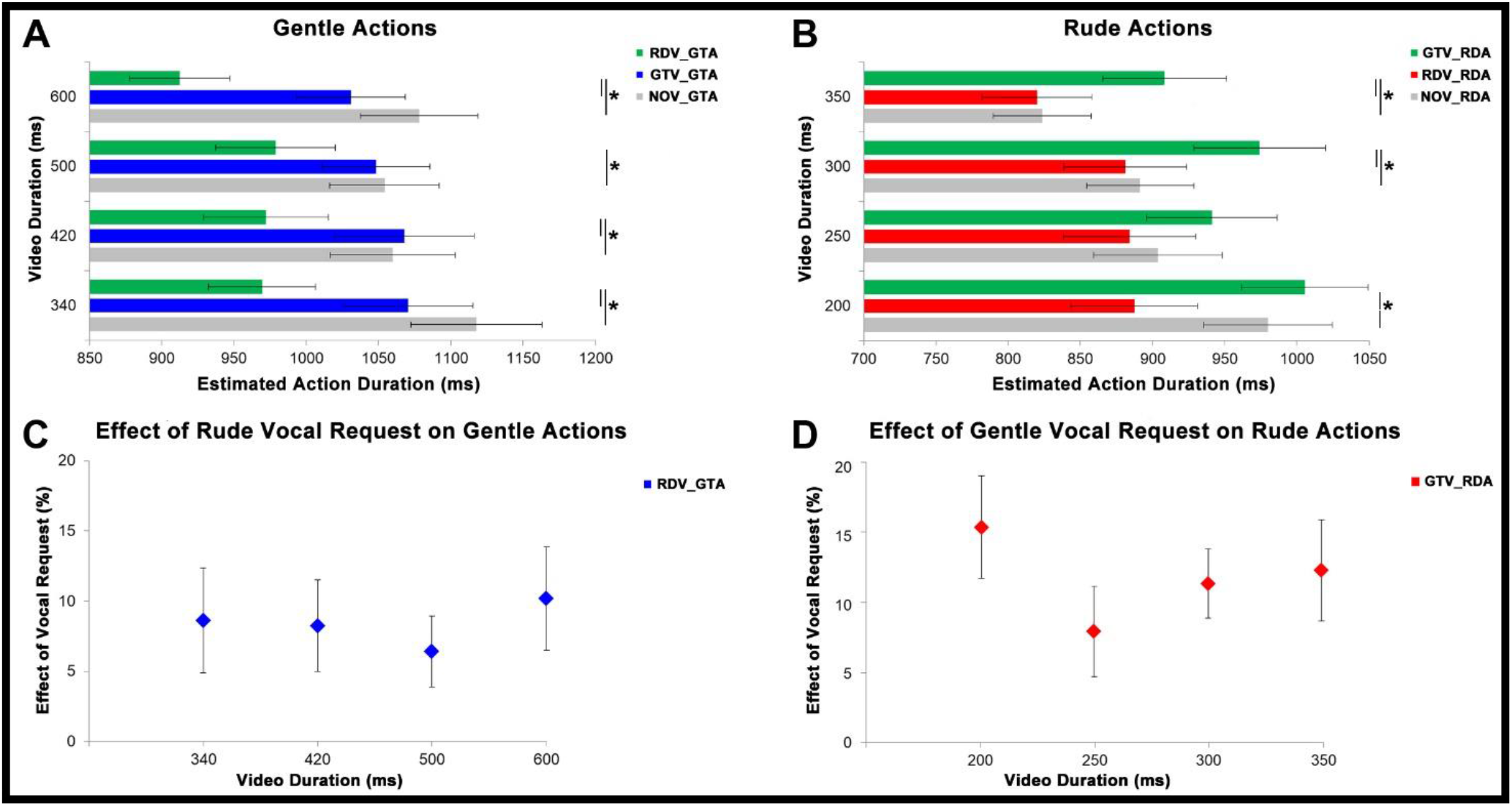
At the top, graphs show results obtained from the analysis of gentle (A) and rude (B) actions estimation. Grey bars refer to the baseline condition (no vocal request: NOV). Green bars refer to incongruent conditions: rude vocal request and gentle action (RDV_GTA), gentle vocal request and rude action (GTV_RDA). Blue bars refer to gentle congruent condition: gentle vocal request and gentle action (GTV_GTA). Red bars refer to rude congruent condition: rude vocal request and rude action (RDV_RDA). At the bottom, graphs show the effect of rude vocal requests on gentle action estimation (C) for 4 different durations (340ms, 420ms, 500ms, 600ms) and the effect of gentle vocal requests on rude action estimation (D) for 4 different durations (200ms, 250ms, 300ms, 350ms).

### 3.2 Second Experiment

In order to evaluate how long the effect of vocal requests conveying different vitality forms last, we analyzed the participants’ responses (action estimation time). Participants’ responses were modelled using two Repeated Measured GLM (general linear models). The significance level was fixed at p = 0.05. Before performing statistical analysis, the sphericity of data was verified (Mauchly’s test, p > 0.05) and the Greenhouse–Geisser correction was applied in case of sphericity violation (p < 0.05). The first GLM model comprised the participant’s response time related to videoclips showing gentle actions after five different time delays (0ms, 400ms, 800ms, 1200ms, 1600ms) in two experimental conditions (congruent: GTV_GTA, incongruent: RDV_GTA). The second GLM model comprised the participant’s response time related to videoclips showing rude actions after five different time delays (0ms, 400ms, 800ms, 1200ms, 1600ms) in two experimental conditions (congruent: RDV_RDA, incongruent: GTV_RDA). Results of the first GLM analysis indicated a significant difference among experimental conditions (F = 16.90, p < 0.001, partial-η^2^ = 0.31, δ = 0.97), time delays (F = 2.33, p < 0.05, partial-η^2^ = 0.05, δ = 0.66) and an interaction between experimental conditions * time delays (F = 2.81, p < 0.05, partial-η^2^ = 0.07, δ = 0.58). For each time delay, post hoc analysis revealed a significant difference among experimental conditions (for 0ms, 400ms and 800ms time delays: GTV_GTA vs RDV_GTA p < 0.001, Bonferroni correction, see Figure 5A). No significant difference was found for time delays of 1200ms and 1600ms. Results of the second GLM analysis indicated a significant difference among experimental conditions (F = 14.20, p < 0.001, partial-η^2^ = 0.27, δ = 0.95) and an interaction between experimental conditions * time delays (F = 3.04, p < 0.01, partial-η^2^ = 0.08, δ = 0.84). For each time delay, post hoc analysis revealed a significant difference among experimental conditions (for 0ms, 400ms and 800ms time delays: RDV_RDA vs GTV_RDA p < 0.001, Bonferroni correction, see Figure 5B). No significant difference was found for time delays of 1200ms and 1600ms. As in the first experiment, in order to quantify the effect of vocal requests on the estimation of action duration for each time delay, we compared values obtained in congruent conditions with those obtained in incongruent conditions (Figure 5CD).

**Figure 5.**
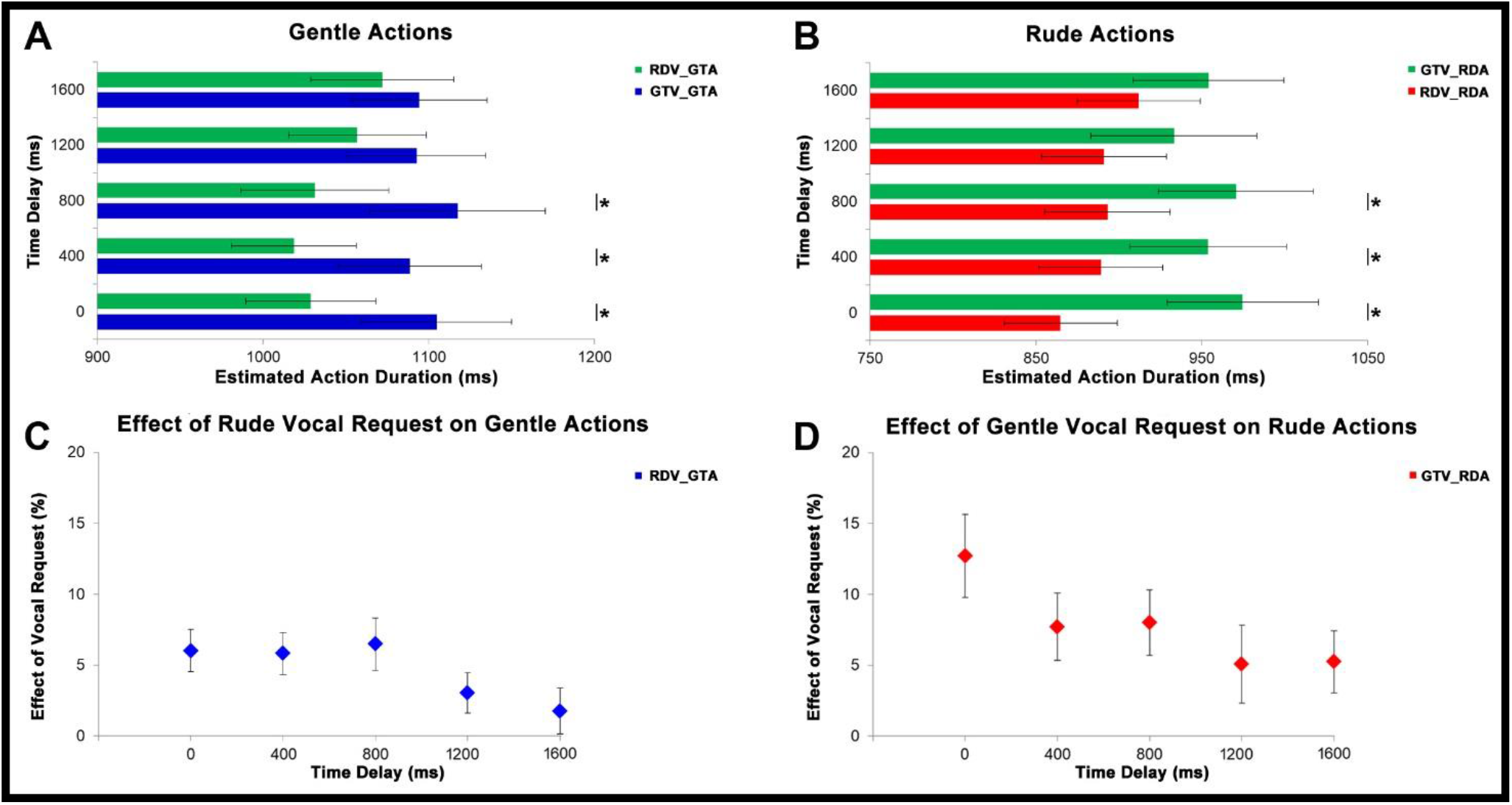
At the top, graphs show results obtained from the analysis of gentle (A) and rude (B) actions estimation. Green bars refer to incongruent conditions: rude vocal request and gentle action (RDV_GTA), gentle vocal request and rude action (GTV_RDA). Blue bars refer to gentle congruent condition: gentle vocal request and gentle action (GTV_GTA). Red bars refer to rude congruent condition: rude vocal request and rude action (RDV_RDA). At the bottom, graphs show the effect of rude vocal requests on gentle action estimation (C) and the effect of gentle vocal requests on rude action estimation (D) for five different time delays (0ms, 400ms, 800ms, 1200ms, 1600ms).

### 3.3 Overall effect of vocal requests on action estimation

Figure 6 shows the overall effect of vocal requests conveying gentle and rude vitality forms on the estimation of action duration. For the first experiment, we averaged values obtained from the comparison between congruent and incongruent conditions for 4 durations (gentle: 340ms, 420ms, 500ms, 600ms; rude: 200ms, 250ms, 300ms, 350ms). For the second experiment, we averaged values obtained from the comparison between congruent and incongruent conditions for the three significant time delays (0ms, 400ms, 800ms). Then a paired sample t-test was carried out to assess possible differences between the effect on gentle action estimation (rude vocal request) and on rude action estimation (gentle vocal request). For both experiments, results showed a significant difference between rude and gentle vitality form (p < 0.05). Specifically, the effect of gentle vocal requests on rude action estimation was greater than the effect of rude vocal requests on gentle action estimation.

**Figure 6:**
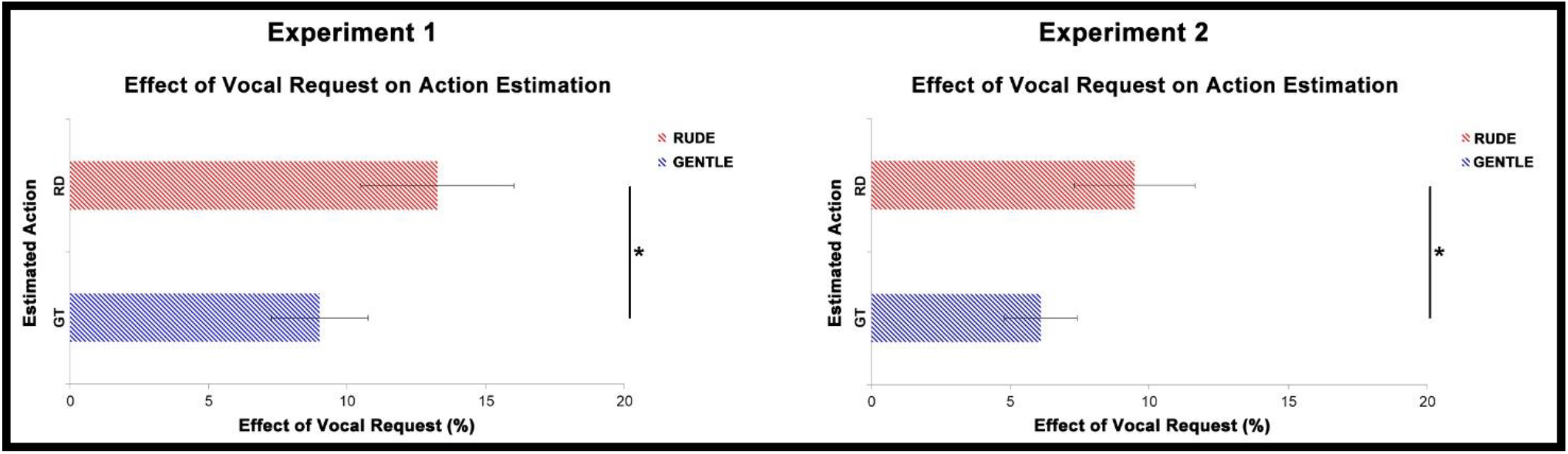
Figure illustrates the overall effect of gentle vocal requests on the estimation of rude (RD, red bars) actions duration and the overall effect of rude vocal requests on the estimation of gentle (GT, blue bars) actions duration. Graphs reported standard errors (SE) and statistical significance.

## 4. Discussion

Observing actions, people may understand two different components: the goal and the form. The goal represents what someone is doing while the form represents the manner in which the action is performed. The form of an action has a strong influence on interaction between humans. According to his affective state, the agent may perform the same gesture in different ways expressing positive or negative attitudes towards the receiver. These subtle aspects of communication have been studied by Daniel Stern, who defined them “vitality forms”.^1^ Although humans continuously convey vitality forms through actions, sounds, speech and touches, to date the impact of vitality forms expressed by an agent on the receiver’s perception has never been investigated. The goals of the present study were two: 1) to evaluate how a positive/negative vocal request may affect the estimation of action duration; 2) to measure the duration of this effect. In the first experiment, the analysis of the participants’ ability to estimate the end of the action showed that listening to a gentle vocal request (“give me”) affected the perception of an action (passing an object), subsequently presented, increasing its duration. In contrast, listening to a rude vocal request decreased the duration of the same action. Specifically, in the incongruent condition, when participants listened to a rude voice and then observed the initial part of a gentle action, anticipated its end. On contrary, if they listened to a gentle voice and then observed the initial part of a rude action, they estimated the action as lasting longer. In the second experiment, results showed that this contagion effect lasted for 800ms and then started to decay. It is plausible to hypothesize that this influence may be ascribed to a potential arousal effect. In order to exclude this possibility, we carried out a further analysis showing that, in both experiments, the effect of gentle vocal request on rude action estimation was significantly greater than the effect of rude vocal request on gentle action estimation. This suggests that the influence of vitality forms on the action estimation was not merely due to an arousal effect because it was easier to slow down the participants’ response than speed it up. It is important to point out that we considered a vocal request consisting in one simple imperative action verb “give me”, pronounced by male or female actors in a rude or gentle way. This limitation may be overcome in the future by reproducing a more realistic dialogue conveying vitality forms. It is plausible that in this way the effect found in the present study may have a longer duration overtime, affecting participants’ behavior in a stronger way.

Our study represents the first demonstration that, observing a small part of a goal directed action, besides the goal, the observer is also able to understand the vitality form of the action. It is plausible that, during the task, participants may have ‘read’ the partial kinematic information of the action and remapped it on their own motor repertoire. This perception-action remapping would have allowed them to simulate internally the vitality forms of action.^11^ This hypothesis is corroborated by several fMRI studies^3-9^ showing that the perception and expression of vitality forms produce the activity of the same neural correlates. Indeed, the perception of action, speech, touches conveying vitality forms, activate the dorso-central insula, more specifically the middle and posterior insula short gyri. Interestingly, the same insular sector is active also during the expression and imaging of actions and speech expressing the same vitality forms (rude, gentle). The authors hypothesized that the dorso-central insula is the key region encoding vitality forms and it is also endowed with a mirror mechanism that makes possible the decoding of the vitality forms of others. Differently from the mirror mechanism located in the parietal and frontal areas, specific for the action goal understanding, ^12-19^ the mirror mechanism located in the insula might allow one to express their own mood/attitude and to understand those of others.^19^

In line with our findings concerning the influence of vitality forms expressed by the agent on the receiver, Di Cesare and colleagues (2017) in a kinematic study showed that, interacting with a virtual agent requiring to take or give an object through different modalities (visual, auditory, mixed), affected some kinematic parameters of participants’ motor responses.^10^ In particular, when participants perceived a rude request they interacted with the object with a larger trajectory and a higher velocity. In contrast, perceiving a gentle request produced a kind interaction with the object, corresponding to a smaller trajectory and a lower velocity. All these findings are corroborated by a very recent study carried out by Lombardi et al. (2021).^20^ More specifically, in a psychophysical study the authors showed that a vocal request expressed rudely or gently by a male actor and female actress, influenced both the perception and execution of actions. Interestingly, the same authors found that the same effect was also present when participants were stimulated with rude or gentle physical requests. Taken together, all these findings suggest how vitality forms conveyed by actions, speech and tactile information can affect people behavior.

The ability to express and understand vitality forms takes a fundamental role in successfully interacting with others. Di Cesare and colleagues showed how specific types of vitality forms as well as changes in vitality forms are particularly difficult to perceive for children with autism spectrum disorder (ASD), suggesting that while subtle expression of these forms of communication may be easily readable for typically developing children (TD), those with ASD may not perceive it.^21^ In a recent kinematic study, Casartelli et. al found that ASD children performed gentle and rude actions differently, as indicated by kinematic parameters (i.e. peak velocity, peak acceleration) but express them in a way that is different from those of typically developing children (TD), suggesting that their difficulties in action understanding may be related to the different way through which they motorically express their own actions.^22^ Taking advantage of actions recorded in this kinematic study, the same authors carried out another study, to assess whether neurotypical adults understand the form of action performed by ASD children.^23^ Particularly, neurotypical adults were presented with two videos showing two different types of actions (placing and throwing) performed with rude or gentle vitality forms by ASD children and TD children. Results showed that they were remarkably inaccurate in identifying vitality forms expressed by ASD children. Since vitality forms are a fundamental aspect of social communication, these findings open the way to a better investigation of the bidirectional difficulties for both ASD and neurotypical people in interacting with each other.

In conclusion, our study provides three main findings. First, we demonstrated that vitality forms conveyed by vocal requests influence the action perception of participants. Second, this contagion effect lasts 800ms and then starts to decay. Finally, we provide first evidence that, observing a goal directed action, besides the goal, the observer is able to internally simulate the vitality forms of that action.

## Acknowledgements

This work has been supported by a Starting Grant from the European Research Council (ERC) under the European Union’s Horizon 2020 research and innovation programme. G.A. No 804388, wHiSPER

## Author Contributions

GDC designed the research; GDC and SA performed the experiment; GDC, AP analyzed the data; GL contributed with material and consulting for the development of the project; GDC and AS wrote the first draft, all authors read the manuscript and contributed to its final form.

## Notes

### Competing Interest Statement

The authors have declared no competing interest.

